# Temporal expectation modulates the neural dynamics of delayed responses to working memory representations

**DOI:** 10.1101/2020.06.08.137539

**Authors:** Fang-Wen Chen, Chun-Hui Li, Bo-Cheng Kuo

**Author notes:** Corresponding Author: Bo-Cheng Kuo, Ph.D., Department of Psychology, National Taiwan University, No. 1, Sec. 4, Roosevelt Road, 10617, Taipei, Taiwan.

## Abstract

Temporal expectation can induce anticipatory attention to enhance perceptual processing and optimise behaviours. However, it remains unexplored whether temporal expectation benefits delayed responses to information maintained in working memory (WM) and the underlying neural dynamics that support the benefits. Here we addressed these issues in behavioural and electroencephalography (EEG) experiments. Participants performed a rotation discrimination WM task. Temporal expectation was manipulated by varying the predictability for when a test probe would occur after a delay interval. We first demonstrated that temporal predictability benefited WM performance. Our EEG results showed decreased alpha power over the left posterior brain regions during the delay interval for high than low predictability trials. Importantly, we found earlier enhancement of the lateralised alpha power and the lateralised readiness potential for high than low predictability trials when WM contents were accessed for responses during the probe period. Finally, we found concurrent selection and preparation for responses when the probe onset was predictable but motor preparation lagged behind response selection when the probe onset was more variable. Together, this study provides new insights into the proactive, dynamic nature of the anticipatory deployment of attention in guiding WM maintenance and delayed responses.

## Introduction

The ability to anticipate upcoming events that are relevant to the task at hand allows individuals to optimise behaviours. Knowing when the relevant events are going to occur particularly enables changes in anticipatory states that confer behavioural advantages with faster or more accurate responses. Many studies have focused on the roles of top-down anticipatory or preparatory attention in perception (Battistoni, Stein, & Peelen, 2017; Nobre & van Ede, 2017; Summerfield & Egner, 2009), isolating distinct processes that enable us to prepare for the relevant stimuli and act upon them. However, an event may occur after a short period of time that requires individuals to select an appropriate response to information maintained in working memory (WM) (D’Esposito & Postle, 2015; Gazzaley & Nobre, 2012; Myers, Stokes, & Nobre, 2017; Stokes, 2015). It remains unclear whether temporal expectation built from the predictability for the onset of test probe could benefit behavioural performance and modulated neural activity associated with delayed responses. In this study, we addressed these issues in behavioural and electroencephalography (EEG) experiments.

A substantial body of research has explored the mechanisms of neural dynamics that underlie anticipatory modulation on upcoming sensory processing. Studies measuring oscillatory activity showed that a spatially specific modulation of cortical excitability in the alpha band (9-13 Hz) over the posterior brain regions reflects orienting of attention in space before the presentation of visual stimuli (Kelly, Lalor, Reilly, & Foxe, 2006; Sauseng et al., 2005; Thut, Nietzel, Brandt, & Pascual-Leone, 2006; Worden, Foxe, Wang, & Simposon, 2000). These findings demonstrated the lateralised alpha effect during the cue-target interval, with a decrease in activity in the contralateral hemisphere but an increase in activity in the ipsilateral hemisphere to the visual field of anticipated inputs. A very similar profile of alpha lateralisation (also referred to as the mu rhythm) (Pfurtscheller, Neuper, & Krausz, 2000) has also been shown in the motor and somatosensory cortices contralateral to the response hand when participants performed tactile discrimination tasks where a visual cue directed attention to their left or right hand (Jones et al., 2010; van Ede, de Lange, Jensen, & Maris, 2011). These findings suggest that such proactive, top-down anticipation indexed by alpha oscillations can optimise both target and response selections when they are relevant to task demands (Jensen & Mazaheri, 2010; Klimesch, Sauseng, & Hanslmayr, 2007).

Growing evidence has revealed that top-down signals may also carry anticipatory codes about the timing of relevant events to guide perception and action (Coull & Nobre, 1998; Ghose & Maunsell, 2002; Nobre & van Ede, 2017; Vangkilde, Coull, & Bundesen, 2012). In this light, temporal expectation could provide critical determinants of attentional prioritisation and selection, thus leading to behavioural improvements if the observers have foreknowledge of when the stimuli will appear. Electrophysiological studies have shown that temporal expectation modulates the strength of alpha oscillations, particularly with a decrease in activity across sensory and motor areas, when anticipating that the relevant stimuli occur at the predicted intervals (van Ede et al., 2011; Zanto et al., 2011) or following a regular rhythmic sequence of stimulations (Rohenkohl & Nobre, 2011). The degree of alpha-power attenuation can be correlated with the enhanced decoding of the target item and distractor resistance if the participants know when the target and distractor are most likely to occur (van Ede, Chekroud, Stokes, & Nobre, 2018).

Recent electrophysiological studies have further revealed that cueing observers to the time when the memory or probe items may occur resulted in changes in alpha activity during the delay interval of WM and concomitantly improve behavioural performance (van Ede, Niklaus, & Nobre, 2017; Wilsch, Henry, Herrmann, Herrmann, & Obleser, 2018; Wilsch, Henry, Herrmann, Maess, & Obleser, 2015). For example, an EEG study manipulated the probability of two items in a memory array to be probed over time in a visual WM task (van Ede et al., 2017). In their study, participants may expect an early or late occurrence of the probe items when they viewed the to-be-remembered items at the beginning of a trial. Their results showed that the dynamic prioritisation of the anticipated item was associated with the temporally specific decrease in contralateral alpha activity before the onset of the test probe. Similar results were reported in auditory WM tasks measured with magnetoencephalography (MEG). Temporal expectation led to a reduction in the cognitive load and a decrease in alpha activity during the retention of syllables in ongoing noise if the observers knew when to listen to the onset of to-be-remembered syllables (Wilsch et al., 2015). However, this was not true when participants were unable to expect the syllables onset during the encoding phase. Moreover, temporal expectation also counteracted the decay of WM traces for pure-tone sequences (Wilsch et al., 2018). When the onset time of the pure-tone sequence was highly predictable for encoding, the WM decline was attenuated along with a decrease in alpha activity over delay durations.

While the beneficial effects of temporal expectation on perception and WM have been observed, it remains unclear whether temporal expectation can modulate delayed responses to the maintained information in WM. In this study, we tested the influences of temporal expectation on delayed responses to WM representations. Moreover, we investigated the neural dynamics that underlie the influences during the delay and probe periods using EEG. Unlike the studies presenting explicit cues (van Ede et al., 2017; Wilsch et al., 2018; Wilsch et al., 2015), we adopted a new experimental approach by varying the duration of delay intervals for a rotation discrimination WM task. The anticipatory states were induced by trials with high or low temporal predictability regarding when the test probe would occur after a short delay. We expected that the temporal expectation would benefit delayed responses and modulate alpha oscillations during the delay interval. Importantly, we tested the effects of temporal expectation on the timing and the strength of electrophysiological activity associated with the delayed responses when the test probe occurred. We expected that the high predictability trials would allow efficient selection of relevant responses, indicated by the lateralised alpha effect. We also tested the effects of temporal expectation on activating response codes and enhancing motor preparation for the responses, as indicated by the lateralised readiness potential (LRP). Together, this study highlights the proactive nature of anticipatory attention induced by temporal expectation in guiding WM maintenance and delayed responses.

## Materials and Methods

Two experiments were conducted, including a behavioural experiment (Experiment 1) and an EEG experiment (Experiment 2). Sixteen healthy volunteers were recruited for each experiment; the behavioural experiment included 7 females (age range 20-24 years, mean age = 20.81 years), and the EEG experiment included 7 females (age range 20-24 years, mean age = 21.00 years). All participants were right-handed, according to the Edinburgh handedness inventory (Oldfield, 1971). They had normal or corrected-to-normal visual acuity, provided written informed consent prior to the study and were financially reimbursed for their time. All experimental methods and procedures were reviewed and approved by the Research Ethics Committee of National Taiwan University.

### Experiment 1

#### Stimuli and apparatus

The stimuli were black and white circular gratings (radius 1.5°) and were presented at the centre of the screen. They were displayed on a 17-inch cathode ray tube (CRT) monitor with a refresh rate of 60 Hz and a resolution of 1024 × 768 pixels. A homogenous grey background was used throughout the experiment. Stimuli were presented with Presentation software (Neurobehavioral Systems, Albany, NY).

#### Task Design

Participants performed a rotation discrimination WM task. They first viewed and remembered the orientation of a sample grating for 200 ms at the centre of the screen. After a varied delay interval, a probe grating appeared for 200 ms, and participants were instructed to indicate which way the probe grating had been rotated compared to the sample grating, using either their right or left thumb, corresponding to clockwise (50% of trials) and counterclockwise (50% of trials) responses, respectively. The sample gratings were presented using one of eight different orientations (11.25 to 168.75 degrees in steps of 22.5). Differences in clockwise and counterclockwise rotations between the sample and the probe gratings were manipulated according to three levels: 5°, 10° and 22.5°. Temporal expectation was manipulated according to the varying duration of the delay interval (i.e., temporal predictability) between the sample grating and the probe grating in separate blocks. On high predictability trials, the delay intervals varied randomly between 1300 and 1700 ms (mean = 1500 ms, uniform distribution). On low predictability trials, the delay intervals varied randomly between 800 and 2200 ms (mean = 1500 ms, uniform distribution). We also included a control condition with a 1500-ms fixed delay interval. The delays with a fixed interval, high and low predictability were presented in a blocked design, and the manipulation of the temporal expectation was never explicitly indicated to the participants. The timing of the probe grating appearance was fully predicted in the control condition and highly predicted in the high predictability condition. It was not possible for participants to predict the onset of probe grating in the low predictability condition. The inter-trial interval, including the 1000 ms response interval, varied randomly between 2000 ms and 3000 ms. The experiment followed a 3 (temporal predictability: fixed, high, low) x 3 (angle difference: 5°, 10° and 22.5°) x 2 (response type: clockwise, counterclockwise) within-subjects factorial design. The task is illustrated in Figure 1A.

**Figure 1.**
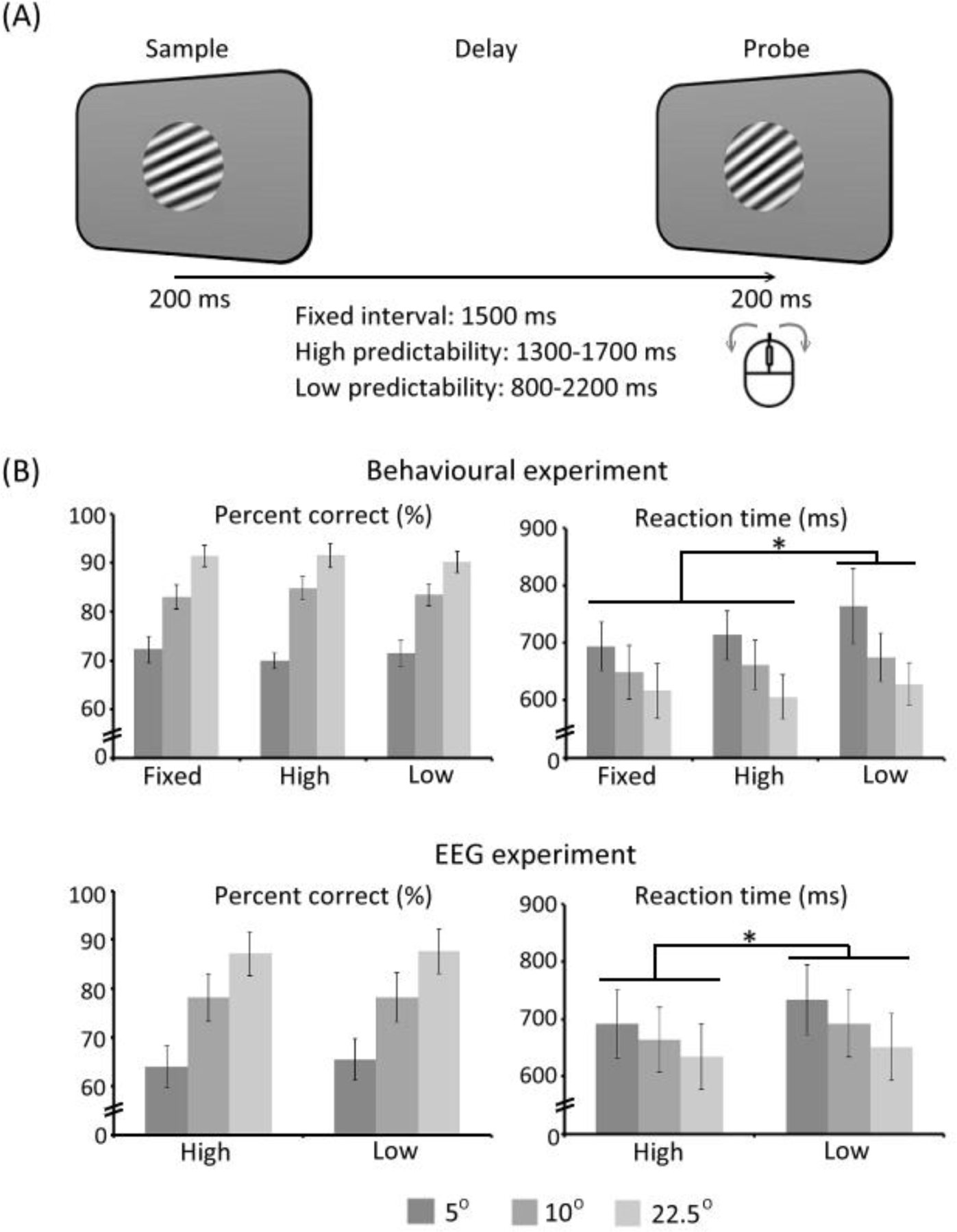
(A) Schematic illustration of the rotation discrimination task. Participants were instructed to indicate which way the probe grating had been rotated compared to the sample grating, using either their right or left thumb, corresponding to clockwise and counterclockwise responses. Temporal expectation was manipulated according to the varying duration of the delay interval between the sample grating and the probe grating. The delay intervals varied randomly between 1300 and 1700 ms for high predictability trials and randomly between 800 and 2200 ms for low predictability trials. A control condition with a 1500-ms fixed delay interval was also included. Differences in clockwise and counterclockwise rotations between the sample and the probe gratings were manipulated according to three levels: 5°, 10° and 22.5°. (B) Behavioural results including mean accuracy (%) and response time (RT, ms) for the behavioural experiment and the EEG experiment. Error bars represent the standard errors of the means.

#### Experimental procedure

Participants were comfortably seated in a dimly illuminated room. Their heads were positioned 60 cm from the CRT monitor using a chinrest. Prior to the formal experiment, the participants were given written, as well as verbal, instructions about the task requirements. Participants first completed a practice block of 8 trials (with a fixed 1500 ms interval) to ensure that they could perform the task as instructed. In the formal experiment, participants performed 9 blocks of 48 trials (three blocks for each temporal predictability condition), yielding 432 trials in total. Response types and angle differences were randomised across trials within a block. The order of the blocks was controlled across participants in a Latin Square design. A fixation marker was displayed at the centre of the screen throughout the whole experiment. Participants were instructed to maintain fixation on a small fixation marker at the centre of the screen during the experimental trials and to respond as quickly and accurately as possible. Participants responded using both thumbs by pressing the mouse buttons. The total experimental time for each subject was approximately 60 minutes.

#### Behavioural analysis

The behavioural measures, including reaction times (RTs) and accuracy, were analysed by 3 (temporal predictability) x 3 (angle difference) two-way repeated-measures analysis of variance (ANOVA). For the RT analysis, only the correct trials were included.

### Experiment 2

The stimuli and design of Experiment 2 (EEG experiment) were identical to those of Experiment 1 (behavioural experiment) with the following exceptions.

#### Design

In the EEG experiment, we only included high and low predictability conditions. The experiment followed a 2 (temporal predictability: high, low) x 3 (angle difference: 5°, 10° and 22.5°) x 2 (response type: clockwise, counterclockwise) within-subjects factorial design.

#### Experimental procedure

The formal experiment consisted of 8 blocks with 48 trials in each block. There were 384 trials in total (192 trials for each predictability condition). Participants were also instructed to minimise their eye movements and blinks within a trial. A fixation point was presented in the centre of the screen throughout the experiment. The total experiment time for each participant was approximately 90 minutes.

#### Behavioural analysis

The RTs and accuracy were each analysed by 2 (temporal predictability) x 3 (angle difference) two-way repeated-measures ANOVA. Only correct responses were included in the RT analysis.

#### EEG acquisition

The EEG was recorded using a NuAmp amplifier (Neuroscan Inc.) with an elastic cap with 37 Ag/AgCl electrodes positioned following the 10-20 international system. The montage arrangement consisted of six midline sites (FZ, FCZ, CZ, CPZ, PZ, and OZ) and twelve sites on each hemisphere (FP1/FP2, F3/F4, F7/F8, FC3/FC4, FT7/FT8, C3/C4, T3/T4, CP3/CP4, TP7/TP8, P3/P4, T5/T6, and O1/O2). Electrodes placed on the supraorbital and infraorbital ridges of the left eye tracked vertical eye movements [vertical electro-oculogram (VEOG)], and horizontal eye movements were recorded using electrodes placed on the outer canthi of the left and right eyes [horizontal electro-oculogram (HEOG)]. Ground (AFz, positioned between FPz and Fz) and reference (A1 and A2 mastoid sites, with A2 serving as the active online reference) electrodes were included throughout the EEG recording. Electrode impedances were kept below 5 KΩ. Continuous brain activity at each electrode site was sampled by 1 ms (1000-Hz analogue-to-digital sampling rate). The activity was filtered with a low-pass filter (300 Hz), and a high-pass filter was not used. Stimulus presentation codes for each event were sent to the EEG acquisition computer via a parallel port from the stimulus presentation computer to indicate the type and exact time of the presentation for each event.

#### EEG preprocessing

For the offline EEG analysis, EEG signals were re-referenced to the algebraic average of the right and left mastoids (A1 and A2). Differences between the voltages at electrodes placed on the side of each eye (HEOG) and above and below the left eye (VEOG) were computed to derive bipolar EOG signals. The continuous EEG data were then segmented into epochs starting 500 ms before and ending 1000 ms after the onset of the sample grating and probe grating. These long epochs prevented windowing artifacts in the time-frequency analysis. The EEG epochs were baseline-corrected, with a 100-ms pre-grating period. Signals containing excessive noise or a steep drift (+/-100 µV) at any electrode were rejected. Epochs between 300 ms before and 800 ms after the grating onset with eye-movement artefacts, including blinks and saccades, were rejected. Blinks were identified as large deflections (+/-60 µV) in the HEOG or VEOG electrodes. Visual inspection was then carried out to confirm the appropriate removal of artefacts and identify residual saccades or eye movements (e.g., boxcar-shaped voltage deflection) in individual HEOG traces (+/- 10-20 µV). Trials with incorrect behavioural responses were also excluded from further analyses. To ensure a sufficient signal-to-noise ratio, a lower limit of 50 artefact-free trials per condition per subject was set.

#### EEG time-frequency power analysis

The offline EEG analyses were performed using SPM12 software (Wellcome Trust Centre for Neuroimaging, University College London, UK) and the Fieldtrip toolbox (Oostenveld, Fries, Maris, & Schoffelen, 2011) in MATLAB (MathWorks) complemented by in-house MATLAB scripts. Epochs of EEG signals were converted into time-frequency decomposition through a short-time Fourier transform of Hanning-tapered data. This transformation was applied to every electrode, trial, and participant across frequencies from 1 Hz to 30 Hz in a 1-Hz step using a fixed 300-ms sliding time window that was advanced over the data in 10-ms steps.

We first tested whether temporal expectation could modulate alpha oscillations (9-13 Hz) during the delay period. The time-frequency power spectrograms were averaged across trials according to the onset of the sample grating for each temporal predictability condition for each participant. Next, we tested whether temporal expectation could modulate neural activity during the test probe. The lateralisation in alpha power provides a measure to characterise the allocation of attention towards the internal selection of responses when the responses involve a choice between the two hands with a greater decreased power over the centre-posterior electrodes contralateral to the response hand (Jones et al., 2010; van Ede et al., 2011; Volberg & Thomaschke, 2017). We calculated the lateralised alpha power for each temporal predictability condition during the probe period. The time-frequency power spectrograms from trials according to the onset of the probe grating containing clockwise responses using the right hand and those containing counterclockwise responses using the left hand were combined by an averaging procedure that preserved the electrode location relative to the required response hand (contralateral or ipsilateral to the response hand) for each temporal predictability for each participant. The power estimates were then log-transformed and rescaled to the baseline relative to -300 ms to -100 ms preceding the grating onset using the LogR method in SPM (log10(p/p_b)) (p: power of the trial; p_b: power in the baseline).

All of the statistical analyses for the time-frequency data were computed using a cluster-based nonparametric permutation method (Maris & Oostenveld, 2007) across participants. This method calculates the size of significant clusters using consecutive *t*-tests that are significant (*p* < 0.05) across neighbouring electrodes, time points, or both, averaging across the frequency band of interest, to control for the problem of multiple comparisons, without making strong prior assumptions, for each participant. Values that exceeded the predefined threshold for *t* tests comparing differences from zero were selected. The selected time points were clustered based on spatial and temporal adjacency. The sums of the *t* values within contiguous clusters were then used for the cluster-based statistics. For the delay period analyses, we tested the significant differences in power between high and low temporal predictability after the sample grating onset. For the probe period analyses, we compared the significant differences in power between the contralateral and ipsilateral sides with respect to the response hand after the probe grating onset for each temporal predictability. We tested the statistical significance using dependent sample *t*-tests by calculating Monte Carlo *p* values using 1000 random permutations, in which two conditions were shuffled to generate the null distribution of the clustered test statistics that would be achieved by chance. Finally, the corrected *p* value was calculated by comparing the values of the observed cluster-level *t* statistics against the null distribution across permutations. The cluster was treated as significant if its size was unlikely to have occurred by chance (*p* < 0.05, two-tailed).

#### Event-related potential (ERP) analysis

To test whether temporal expectation can modulate electrophysiological activity leading to the preparation of motor responses, we also calculated the LRP (Coles, 1989; Gratton, Coles, Sirevaag, Eriksen, & Donchin, 1988; Kutas & Donchin, 1980) for each temporal predictability condition during the probe period. The LRP provides a suitable measure to characterise the effect of motor preparation when the responses involve a choice between the two hands, with greater negativity over the frontal-central electrodes contralateral to the response hand. In our study, the LRP was derived from the onset of the probe grating and then averaged according to temporal predictability and response hand for each participant. The epochs from trials containing responses made by the right hand (clockwise response) and from trials containing responses made by the left hand (counterclockwise response) were combined by an averaging procedure that preserved the electrode location relative to the response hand (contralateral or ipsilateral to the response hand). The mean amplitudes of the LRP were measured at the FC3/C3 and FC4/C4 electrodes sensitive to motor and premotor activity from 300 ms to 700 ms relative to the probe onset by computing the difference in the mean amplitude between the contralateral and ipsilateral electrodes with respect to the response hand (Coles, 1989; Miller, Patterson, & Ulrich, 1998; MÜller-Gethmann, Ulrich, & Rinkenauer, 2003). The LRP was analysed by 2 (temporal predictability) x 2 (electrode side: contralateral, ipsilateral) two-way repeated-measures ANOVA.

#### Time-course analyses for the probe-related activity

To further explore the influences of temporal expectation on the time courses of the alpha lateralisation and the LRP during the probe period, we divided the period of 300-700 ms after the probe grating onset into four time windows: 300-400 ms, 400-500 ms, 500-600 ms, and 600-700 ms. We extracted the mean power estimates of alpha activity and mean amplitudes of the LRP from the high and low predictability conditions, respectively, for each of the four time windows from the electrodes of interest (alpha: CP3/4 and P3/4; LRP: FC3/4 and C3/4) for each participant. The alpha power estimates and the LRP amplitudes were then subjected to a three-way repeated-measures ANOVA, testing the effects of temporal predictability, time window (300-400 ms, 400-500 ms, 500-600 ms, 600-700 ms) and electrode side. Only effects including the electrode side, reflecting differential activity observed in the contralateral versus the ipsilateral electrodes, were of interest. For the statistical analyses of EEG/ERP data as well as behavioural data, a Greenhouse-Geisser correction was used if sphericity was violated.

Moreover, we tested the temporal relationship between the time courses of lateralised alpha power and the LRP using a cross-correlation analysis for each temporal predictability condition. We first subtracted the power/amplitude of the alpha activity/LRP at the ipsilateral side from the contralateral side relative to the response hand for each condition for each participant and then calculated the cross-correlational coefficients (*r*) between the two time courses from 300 ms to 700 ms using the *xcov* function in MATLAB. We set the range of time lag from -400 ms to 400 ms, shifting the time course of the LRP in a 10-ms step on the basis of the time course of the lateralised alpha power. The cross-correlation coefficient varies from -1 to 1, where 1 (−1) indicates a positive (negative) correlation between two signals and 0 indicates no correlation. The maximal coefficient at a negative or positive time lag indicates the temporal difference in the correlation between the time courses of lateralised alpha activity and the LRP. A negative time lag reflects that the LRP precedes the lateralised alpha activity. A positive time lag reflects that lateralised alpha activity precedes the LRP. The maximal coefficient at a zero lag indicates that there is no temporal difference between the two time courses. The cross-correlation coefficients were transformed into *z* values using Fisher’s *z* transform for each participant. We used one sample *t*-tests (*p* < 0.05, one-tailed) to compare Fisher’s *z* scores against zero (i.e., no correlation) for each time lag for each predictability condition. We also extracted the time lags on the basis of the maximal cross-correlation coefficient for each predictability condition for each participant and evaluated the time lags using a Jackknife method (Miller et al., 1998). We iteratively removed the data for one participant from the data for all participants, resulting in 16 new datasets. This leave-one-participant-out approach allows us to obtain a Jackknife estimate of the reliability of the time lags and compare the difference in the time lags between high and low predictability conditions using a paired *t*-test (*p* < 0.05, two-tailed).

#### Relationship between the probe-related activity and RTs

Finally, we conducted a model-based analysis to test for the relationship between the probe-related activity (the lateralised alpha power and the LRP) and the RTs at two levels. At the individual level, we calculated the lateralised alpha power/LRP by subtracting the power/amplitude at the ipsilateral side from the contralateral side relative to the response hand for each trial for each participant. We then fitted a general linear model (GLM) to the *z*-transformed lateralised alpha power/LRP using the *z*-transformed RT as a regressor at each time point for each electrode of interest on a trial-by-trial basis. The GLM was fitted for each of the temporal predictability conditions separately. This was repeated for each participant’s dataset. Fitted data for each individual (beta estimates) were then entered into the group-level analysis. We used one sample *t*-tests to examine a significant relationship between the LRP/lateralised alpha power and RTs (*p* < 0.05, two-tailed). In essence, this analysis tests whether the variability in the RT is significantly predicted by the trial-wise variability in the magnitude of the lateralised alpha power/LRP.

## Results

### Behavioural results

The behavioural results are illustrated in Figure 1B. In the behavioural experiment, accuracy data showed a significant main effect of angle difference [*F*(2,15) = 115.08, *p* < 0.001], indicating better performance for a 22.5° difference (91.06 ± 2.13 %) compared to 5° (71.23 ± 1.94 %) and 10° (83.77 ± 2.08 %) differences and for a 10° difference compared to a 5° difference. RT data revealed a significant main effect of temporal predictability [*F*(1.41, 21.11) = 5.75, *p* = 0.017] and a significant main effect of the angle difference [*F*(2, 30) = 27.20, *p* < 0.001]. These results indicated faster RTs for the trials with a fixed interval (653.47 ± 44.53 ms) and high predictability (660.80 ± 40.08 ms) compared to the trials with low predictability (689.17 ± 45.96 ms) and faster RTs for a 22.5° difference (616.97 ± 40.09 ms) compared to 5° (724.32 ± 48.32 ms) and 10° (662.15 ± 43.00 ms) differences and for a 10° difference compared to a 5° difference. No other effect was significant.

In the EEG experiment, accuracy data replicated that from the behavioural experiment and showed a significant main effect of angle difference [*F*(2,30) = 151.18, *p* < 0.001], indicating better performance for a 22.5° difference (91.91 ± 1.31 %) compared to 5° (67.69 ± 1.78 %) and 10° (82.77 ± 1.78 %) differences and for a 10° difference compared to a 5° difference. RT data also replicated the effects observed in the behavioural experiment and showed a significant main effect of temporal predictability [*F*(1, 15) = 11.79, *p* = 0.004] and a significant main effect of angle difference [*F*(1.39, 20.78) = 17.93, *p* < 0.001]. These results indicated faster RTs for the high predictability condition (708.76 ± 37.03 ms) compared to the low predictability condition (740.19 ± 36.66 ms) and for a 22.5° difference (686.51 ± 39.92 ms) compared to 5° (763.94 ± 35.91 ms) and 10° (722.98 ± 35.98 ms) differences and for a 10° difference compared to a 5° difference. We also found a significant interaction between the temporal predictability and the angle difference [*F*(2, 30) = 7.18, *p* = 0.003]. This interaction arose because the effect of temporal expectation with a faster RT for high predictability trials relative to low predictability trials was significant for 5° (*p* < 0.001) and 10° differences (*p* = 0.006).

### EEG time-frequency and time-course results

Time-frequency analyses of EEG data during the delay period were first performed to test whether the power of alpha activity was modulated by the temporal expectation in a way that could change cortical excitability according to the anticipated probes. We found that the high predictability condition induced sustained and decreased power in the alpha band relative to that in the low predictability condition over the left parietal-occipital electrodes from 230 ms to 660 ms after the sample grating onset (corrected *p* = 0.035) (Figure 2).

**Figure 2.**
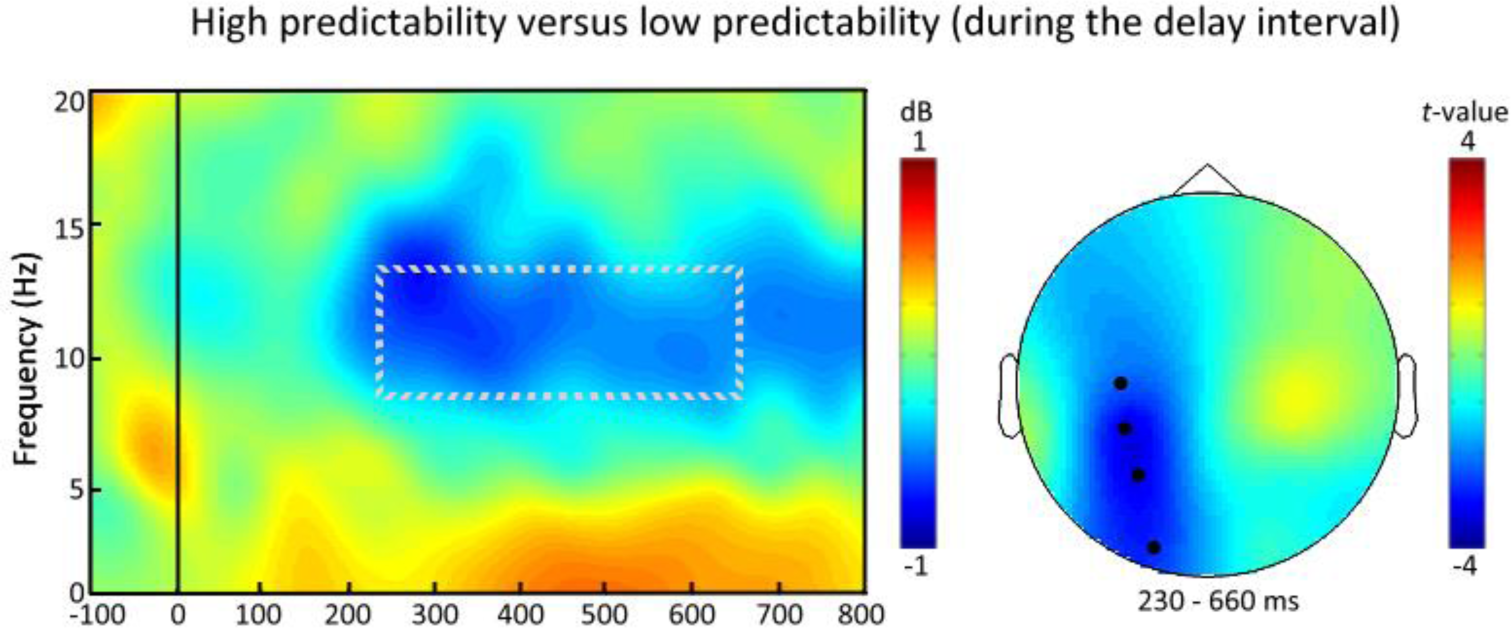
Time-frequency analyses of EEG data during the delay period showed that high predictability trials induced a sustained and decreased power in the alpha band relative to that for low predictability trials over the left parietal-occipital electrodes (C3, CP3, P3 and O1) from 230 ms to 660 ms after the sample grating onset. The time window for the significant effect is indicated by a square (left panel). The topographic map shows the difference in alpha power between the high and low predictability conditions (right panel).

Importantly, we tested whether the temporal expectation can bias the neural responses during the probe period. The analysis of the spectral lateralisation of alpha power relative to the response hand yielded significant effects of attentional selection over centre-posterior electrodes after the probe grating onset. The relative attenuation of alpha power contralateral (versus ipsilateral) to the relevant response hand was observed from 390 ms to 750 ms after the probe grating onset for the high predictability condition (corrected *p* = 0.034). A similar attenuation of alpha lateralisation was observed from 610 ms to 700 ms for the low predictability condition (corrected *p* = 0.01).

The time-course analysis of alpha power during the probe period showed a significant effect of electrode side [*F*(1, 15) = 8.07, *p* = 0.012], suggesting greater alpha attenuation at the contralateral side versus the ipsilateral side to response hand. We also found a significant interaction between temporal predictability and time window [*F*(3, 45) = 2.72, *p* = 0.05] and a significant interaction between time window and electrode side [*F*(3, 45) = 6.69, *p* = 0.001]. We then tested these effects for each time window in detail in a two-way (temporal predictability and electrode side) repeated-measures ANOVA. For the time window of 300 to 400 ms, we found a significant interaction between temporal predictability and electrode side [*F*(1, 15) = 5.83, *p* = 0.03], indicating that the lateralised effect in alpha power was more pronounced for high predictability trials than low predictability trials. We also observed a significant effect of electrode side for the other three time windows [400-500 ms: *F*(1, 15) = 5.54, *p* = 0.03; 500-600 ms: *F*(1, 15) = 8.59, *p* = 0.01; 600-700 ms: *F*(1, 15) = 20.82, *p* < 0.001], which confirmed the decrease in alpha power at the centre-posterior electrodes contralateral to response hand compared to that at the ipsilateral side. Finally, we observed a significant effect of temporal predictability for the time window of 600 to 700 ms [*F*(1, 15) = 4.62, *p* = 0.05], reflecting a greater decrease in alpha power for low predictability trials compared to high predictability trials. These results suggest that temporal expectation can modulate lateralised alpha power during the probe period starting at approximately 300 ms when the probe onset was highly predictable. The time-frequency and time-course results of alpha lateralisation are shown in Figure 3.

**Figure 3.**
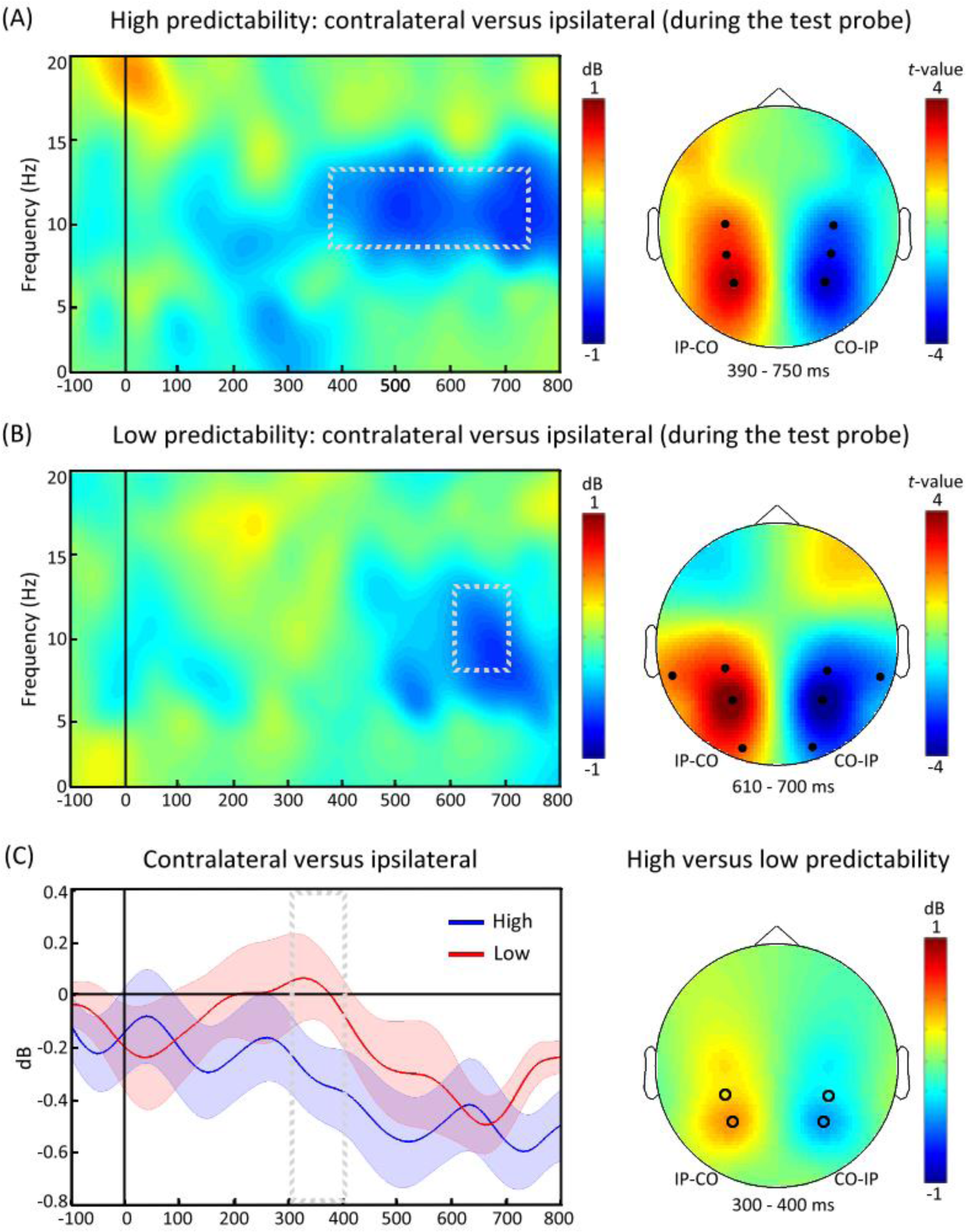
(A) Analysis of the spectral lateralisation of alpha power relative to the response hand yielded significant effects of attentional selection over centre-posterior electrodes (C3/4, CP3/4 and P3/4) after the probe grating onset. The relative attenuation of alpha power contralateral (versus ipsilateral) to the relevant response hand was observed from 390 ms to 750 ms after the probe grating onset for high predictability trials. (B) A similar attenuation of alpha lateralisation was observed from 610 ms to 700 ms over centre-posterior electrodes (TP7/8, CP3/4, P3/4, and O1/2) for low predictability trials. (C) The time-course analysis of alpha power during the probe period showed that the lateralised effect in alpha power corresponding to the response hand was more pronounced for high predictability than for low predictability for the time window of 300 to 400 ms over posterior electrodes (e.g., CP3/4 and P3/4). The differences in power between the ipsilateral and contralateral electrodes are illustrated for high predictability (blue) and low predictability trials (red) (left panel). The shadow represents the standard errors of the means. The time window of interest is indicated by a square (left panel). The topographic map shows the symmetrical relative difference in alpha power (right panel), with increased power over the ipsilateral (IP) scalp and decreased power over the contralateral (CO) scalp. Blue indicates decreased power (CO-IP), and red indicates increased power (IP-CO).

### ERP and time-course results

We also exploited the LRP to test for the influences of temporal expectation on motor preparation during the probe period. Overall, the analysis of the mean amplitude of the LRP from 300 to 700 ms showed a significant effect of electrode side [*F*(1, 15) = 12.73, *p* = 0.003], indicating a more negative waveform at the contralateral electrodes relative to the ipsilateral electrodes to the response hand. A significant effect of time window was observed [*F*(3, 45) = 13.37, *p* < 0.001], suggesting that the LRP amplitude varied over time. The interaction between temporal predictability and electrode side was also significant [*F*(1, 15) = 5.92, *p* = 0.027], suggesting a greater LRP for high predictability trials than that for low predictability trials. Moreover, we tested the LRP effect for each time window in detail in a two-way (temporal predictability and electrode side) repeated-measures ANOVA. We found a significant interaction between temporal predictability and electrode side across three time windows [300-400 ms: *F*(1, 15) = 6.71, *p* = 0.02; 400-500 ms: *F*(1, 15) = 8.23, *p* = 0.011; 500-600 ms: *F*(1, 15) = 4.35, *p* = 0.05] but no such interaction effect for the time window of 600-700 ms (*p* > 0.1). These interactions suggested a greater LRP for the high predictability condition relative to that for the low predictability condition from approximately 300 ms to 600 ms. The LRP effect became comparable between the two predictability conditions at approximately 600 ms. The LRP and time-course results are shown in Figure 4.

**Figure 4.**
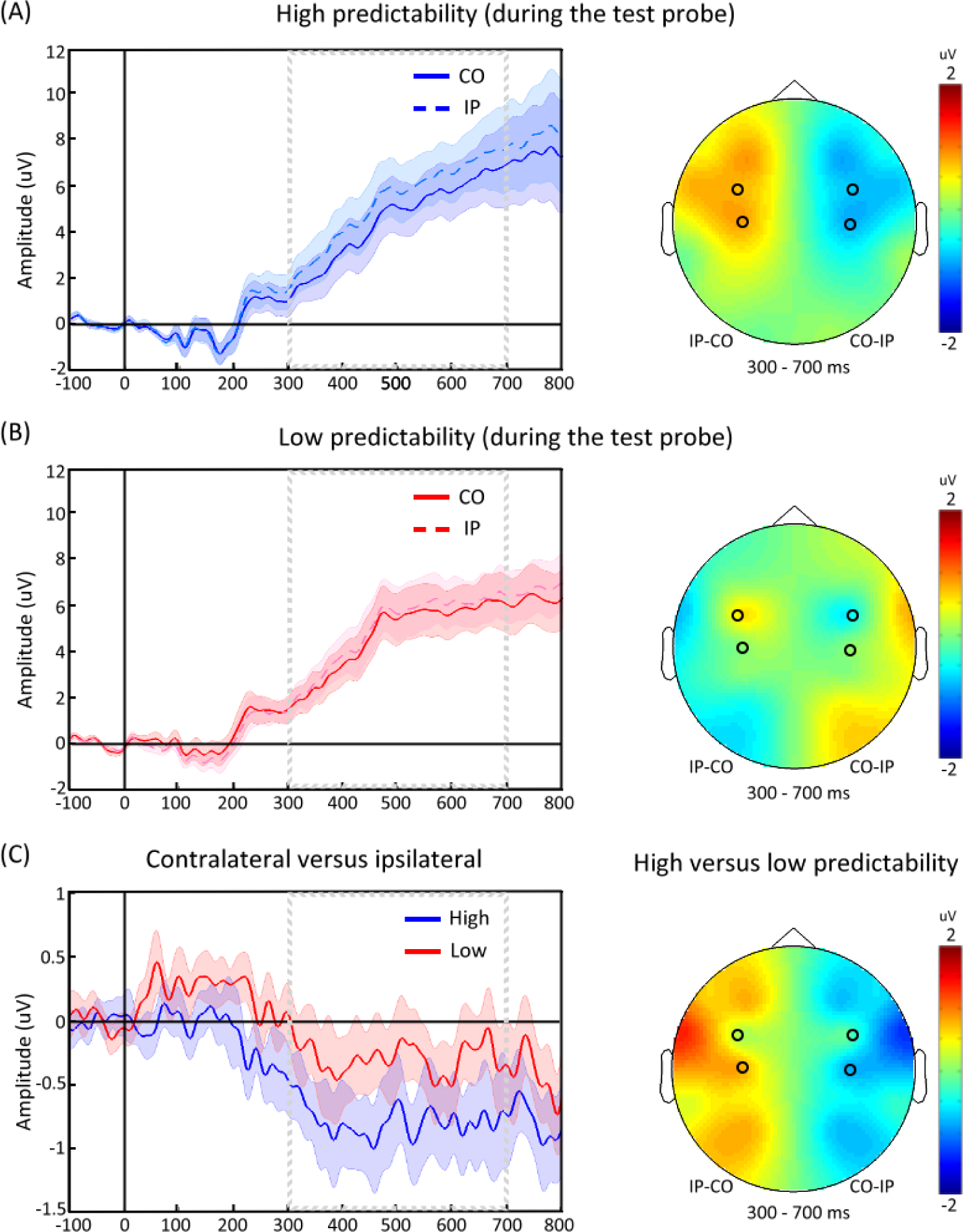
The LRP was measured and analysed between 300 and 700 ms at the electrode pairs (FC3/4 and C3/4) contralateral (CO, waveforms: solid lines) and ipsilateral (IP, waveforms: dashed lines) to the side of the response hand for high predictability (A) and low predictability (B). (C) The voltage differences between the ipsilateral and contralateral sides are shown for high predictability (blue) and low predictability trials (red). The ERP waveforms were low-pass filtered (40 Hz) for illustration purposes. The shadow represents the standard errors of the means. The time window of interest is indicated by a square (left panel). The topographic map shows the symmetrical relative difference in amplitude (right panel). Blue indicates a negative amplitude (CO-IP), and red indicates a positive amplitude (IP-CO).

### Temporal relationship between the LRP and lateralised alpha activity

Next, we calculated cross-correlations between the time courses of lateralised alpha power and LRP for each participant. We found that the lateralised alpha power was significantly correlated with the LRP at a time lag of approximately zero for the high predictability condition [corrected *t*(15) = 3.41, *p* < 0.05] and at positive time lags for the low predictability condition [corrected *t*(15) = 3.63, *p* < 0.05]. We then compared the difference in the time lag between high and low predictability conditions. This analysis showed that the maximal coefficient of cross-correlation between the lateralised alpha power and the LRP occurred earlier for the trials with high predictability (11.88 ± 31.03 ms) relative to those with low predictability (62.5 ± 37.86 ms) [corrected *t*(15) = 3.68, *p* < 0.05] (Figure 5).

**Figure 5.**
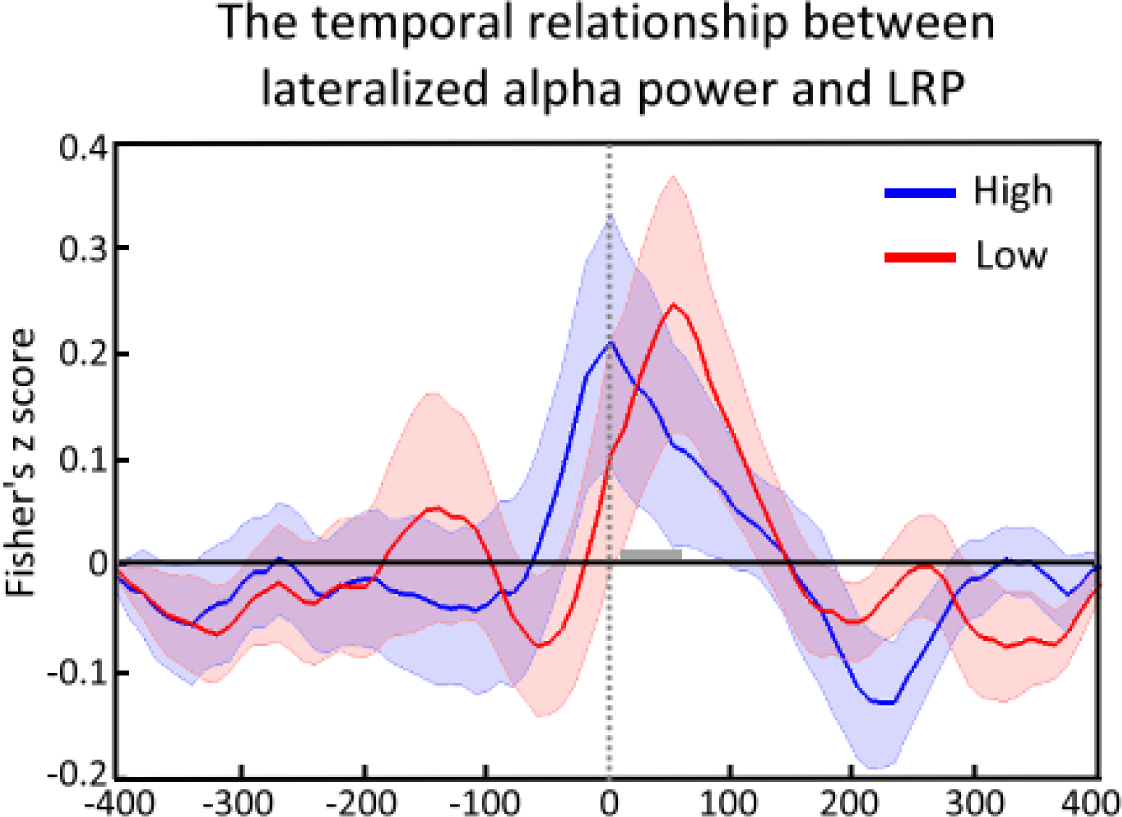
The cross-correlation coefficients between the time courses of lateralised alpha power and the LRP were evaluated for each participant. We set the range of the time lag from - 400 ms to 400 ms, shifting the time course of the LRP in a 10-ms step on the basis of the time course of the lateralised alpha power. We found that the lateralised alpha power was significantly correlated with the LRP at a time lag of approximately zero for high predictability trials (blue) and at positive time lags for low predictability trials (red). We showed that the maximal coefficient of cross-correlation between the lateralised alpha power and the LRP occurred earlier for the trials with high predictability relative to the cross-correlation between those with low predictability (grey bar). The shadow represents the standard errors of the means.

### Relationship between the probe-related activity and RTs

Finally, we conducted a trial-by-trial GLM analysis to test for a relationship between the lateralised alpha power/LRP and the RTs. We showed a significant positive relationship between the LRP and RTs from approximately 350 ms to 490 ms on high predictability trials (corrected *p* = 0.047) and from approximately 350 ms to 520 ms on low predictability trials (corrected *p* = 0.01). These results showed that LRP amplitude was correlated with the RTs for both predictability conditions, suggesting the larger the LRP (more negative) the faster the RT, on that trial for both predictability conditions. No significant relationship between the lateralised alpha power and RTs was observed (*p*s > 0.1).

## Discussion

In this study, we investigated whether temporal expectation can benefit WM performance and modulate the neural dynamics of delayed responses to WM representations across two experiments. By manipulating the temporal predictability regarding when the test probe would occur in a rotation discrimination WM task, we first demonstrated behavioural benefit by temporal predictability. Moreover, our EEG data showed a decrease in alpha power over the left posterior regions during the delay interval for high predictability trials relative to that for low predictability trials. Importantly, we showed earlier enhancement of the lateralised alpha power and the lateralised readiness potential for high than low predictability trials when WM contents were accessed for responses during the probe period. Finally, we found a positive correlation in the time course between the lateralised alpha power and LRP for both predictability conditions. Response selection and motor preparation engaged in parallel when the probe onset was predictable but these two processes engaged in succession when the probe onset was variable. Together, this study provides novel evidence into the proactive, dynamic nature of the anticipatory deployment of attention in guiding WM maintenance and delayed responses.

As the available evidence has revealed that temporal expectation can facilitate the processing of anticipated events if observers have foreknowledge about when those events are going to occur (Coull & Nobre, 1998; Ghose & Maunsell, 2002; Nobre & van Ede, 2017), we showed that temporal expectation had a similar influence on delayed-response performance. Our behavioural data suggested that high predictability trials were effective at inducing anticipatory states with respect to when the probe stimuli will likely appear at particular times. Conversely, delayed responses were less efficient for low predictability trials when the duration of the delay interval was highly variable. These results were consistent with the finding of a recent behavioural study in which cueing participants to a point in time can lead to perceptual improvements at expected time points but impairments at unexpected time points (Denison, Heeger, & Carrasco, 2017). We suggest that the duration of delay interval in our study may serve as a temporal cue and establish expectations about the occurrence of the test probe. Temporal expectation can thus facilitate delayed responses.

Our time-frequency analysis of alpha power during the delay interval showed a significant decrease in alpha power over the left posterior brain regions when the probe onset was highly predictable relative to that when it was highly variable. These results replicated and extended the neural evidence from the recent EEG and MEG studies in WM tasks (van Ede et al., 2017; Wilsch et al., 2018; Wilsch et al., 2015), reflecting the changes in neural states in anticipation of the test probe. More specifically, this hemispheric lateralisation of alpha attenuation from our EEG results is also in agreement with the neuroimaging findings indicating that temporal orienting can preferentially activate the left parietal cortex, especially around the intraparietal sulcus, when participants used temporal cues to predict the time of target onset (Coull, Cotti, & Vidal, 2016; Coull & Nobre, 1998; Davranche, Nazarian, Vidal, & Coull, 2011). Together, our results suggest that the anticipatory state during WM maintenance can be reconfigured by implicitly induced temporal predictability. This top-down anticipatory modulation of WM representations is supported by the changes in cortical excitability in the alpha band in the left posterior brain regions and exerts a proactive influence on WM maintenance.

Importantly, a novel finding from the current study is that the anticipatory states induced by temporal expectation modulated probe-related activity when the WM content was accessed for responses in two aspects. First, our time-frequency results showed a corresponding decrease in alpha power at centre-posterior electrodes contralateral to the required response hand on both high and low predictability trials. The time-course analysis further revealed that a greater effect of alpha lateralisation occurred earlier for high predictability trials than that for low predictability trials. Interestingly, the lateralised alpha effect in the current study is similar to the posterior alpha lateralisation when orienting attention to the location of the upcoming stimulus (Kelly et al., 2006; Rihs, Michel, & Thut, 2009; Sauseng et al., 2005; Thut et al., 2006; Worden et al., 2000) and the somatosensory alpha lateralisation when orienting attention to the response hand (Haegens, Luther, & Jensen, 2012; van Ede et al., 2011). Here, we showed the modulation of alpha lateralisation across somatosensory and parieto-occipital regions when the participants directed attention to the relevant responses according to the comparisons between the tested and maintained information. These results support the notion that alpha lateralisation may reflect a general mechanism that determines the engagement and disengagement of the cortices according to the task relevancy across various sensory modalities and controls the representational states of WM (de Vries, Slagter, & Olivers, 2020; Klimesch, 2012; Kuo, Li, Lin, Hu, & Yeh, 2017; van Ede, 2018; Wallis, Stokes, Cousijn, Woolrich, & Nobre, 2015). We showed that temporal predictability can optimise attentional prioritisation and the selection of responses, thereby bringing alpha lateralisation forward when the probe onset was highly predictable.

Second, our analyses of the probe-locked LRP provided additional evidence in support of the anticipatory influence of temporal expectation on delayed responses. Studies measuring the LRP have shown that temporal expectation exerts a great influence on response preparation by enhancing the speed of both perceptual and motor processing (Faugeras & Naccache, 2016; MÜller-Gethmann et al., 2003; Praamstra & Pope, 2007; Rohenkohl & Nobre, 2011; Volberg & Thomaschke, 2017). For example, a previous EEG study that compared regular rhythmic versus irregular arrhythmic patterns of stimulations showed that rhythm-induced temporal expectation led to both an earlier onset and a larger amplitude of the LRP (Rohenkohl & Nobre, 2011). A recent study presented visual stimuli after a short or long foreperiod and required participants to respond using either the left or the right hand (Volberg & Thomaschke, 2017). Their results showed that the polarity of the LRP can be reversed after the short foreperiod. Consistent with these findings, we showed that the LRP effect was brought forward and enhanced in amplitude for high predictability trials relative to that for low predictability trials. These results reflect that high predictability trials may advance the preparation of motor responses to initiate the appropriate action. Finally, we found a positive relationship between LRP and RTs – the more negative the LRP, the faster the RTs, for both predictability conditions. Since we did not observe a significant correlation between other oscillatory measures and behavioural performance, a possible explanation is that the link between motor preparation and RTs is essential for delayed responses regardless of its temporal predictability.

In sum, our results of probe-related activity suggest that temporal expectation is highly prevalent in delayed responses, favouring hand-specific response activation. While the relevant responses were prioritised and selected, as indicated by the alpha lateralisation, the motor responses can be prepared, as indicated by the LRPs, during the majority of the probe course. Both effects were set to become amplified earlier when the probe onset was predictable compared to those when it was variable. Temporal expectation can thus enhance the selection and preparation of the responses to maintained representations.

While we have highlighted the distinct functions of alpha lateralisation and the LRP during the probe period as the temporal predictability was varied, the temporal relationship between these two signals is unclear. To address this issue, we calculated the cross-correlational coefficients between the time courses of the lateralised alpha power and LRP. Our analyses revealed a positive correlation at time lags of approximately zero for the high predictability condition but at positive time lags for the low predictability condition. When the probe onset was highly predictable, the LRP occurred at approximately the same time as the lateralised alpha effect. When the probe onset was highly variable, the LRP could occur following the lateralised alpha effect (approximately 50 ms later).

We suggest that responding to the test probe may activate dynamic and multiple top-down mechanisms: prioritisation/selection and preparation of relevant motor responses. One mechanism is to direct attention towards the relevant response based on current goals. This process allows for prioritising and selecting relevant responses when comparing the incoming inputs with the mnemonic representations (Hyun, Woodman, Vogel, Hollingworth, & Luck, 2009; Kuo & Astle, 2014; Kuo, Rotshtein, & Yeh, 2011; Myers et al., 2015) according to the predefined task demand (in our case, clockwise or counterclockwise). Another mechanism is to prepare for response initiation. This process allows a person to plan to execute specific motor acts. A recent MEG study provided complementary evidence to highlight this account of dynamic and multiple top-down mechanisms (van Ede, Chekroud, Stokes, & Nobre, 2019). They associated individual memory items with particular responses and tracked the neural dynamics of visual (e.g., item location) and motor (e.g., response hand) selections for WM utilisation. When the mnemonic codes have both visual and motor attributes, the selections of attributes involved neural responses in both the visual and motor areas over highly similar intervals after the probe onset. These two effects also exhibited concurrent activation in the time course, thereby suggesting coactivation of visual and motor selections. We extended their results, focusing on the anticipatory modulations of delayed responses. We showed that temporal expectation can modulate prioritisation/selection and preparation of delayed responses during the probe period. When the probe onset can be predicted, the LRP can be generated as soon as the selection of the response has begun. The relevant response can be prioritised while carrying their motor affordance at the same time, thereby affording efficient actions. In contrast, the LRP may lag behind response selection when the probe onset cannot be predicted. Selection and preparation for the delayed responses may thus engage in succession.

In conclusion, we have isolated multiple processes in WM that benefit from temporal predictability. We provided evidence showing that temporal expectation can reduce temporal uncertainty about the timing of the test probe and facilitate delayed responses. To our knowledge, this is the first evidence suggesting that temporal expectation dynamically influences the strength and timing of probe-related electrophysiological responses, which are reflected by the modulations of lateralised alpha power and LRP, when the WM contents are accessed for delayed responses. Differential influences by temporal expectation on the relationships between the neural markers provide novel evidence for the distinct roles of response selection and preparation in guiding WM responses.

## Acknowledgements

This work was supported by grants from the Ministry of Science and Technology, Taiwan (MOST 104-2628-H-002-002-MY3) and the Ministry of Education, Taiwan (NTU-108L7850) to B.-C. Kuo. F.-W. Chen and C.-H. Li contributed equally to this work. We are grateful to the insightful comments provided by Dr. Yei-Yu Yeh. Correspondence should be addressed to B.-C. Kuo.

## Notes

### Competing Interest Statement

The authors have declared no competing interest.

